# Improved Protocol for Single-Nucleus RNA-sequencing of Frozen Human Bladder Tumor Biopsies

**DOI:** 10.1101/2022.10.14.512220

**Authors:** Sofie S. Schmøkel, Iver Nordentoft, Sia V. Lindskrog, Philippe Lamy, Michael Knudsen, Jørgen Bjerggaard Jensen, Lars Dyrskjøt

## Abstract

This paper provides a laboratory workflow for single-nucleus RNA-sequencing (snRNA-seq) including a protocol for gentle nuclei isolation from fresh frozen tumor biopsies, making it possible to analyze biobanked material. To develop this protocol, we used non-frozen and frozen human bladder tumors and cell lines. We tested different lysis buffers (IgePal and Nuclei EZ) and incubation times in combination with different approaches for tissue and cell dissection; sectioning, semi-automated dissociation, manual dissociation with pestles, and semi-automated dissociation combined with manual dissociation with pestles. Our results showed a combination of IgePal lysis buffer, tissue dissection by sectioning and short incubation time was the best conditions for gentle nuclei isolation applicable for snRNA-seq, and we found limited confounding transcriptomic changes based on the isolation procedure. This protocol makes it possible to analyze biobanked material from patients with well described clinical and histopathological information and known clinical outcomes with snRNA-seq.

## Introduction

The field of single cell genomics has developed rapidly during the last years and allows investigation of tissue at single cell resolution. Single-cell RNA-sequencing (scRNA-seq) has been used to identify novel cell types and cell states^1,2^. Furthermore, the method has also been used to reveal the composition of the tumor ecosystem in e.g. metastatic melanoma, and to detect rare cell subpopulations and unravel intra-tumor heterogeneity^2–6^. However, frequently applied protocols for scRNA-seq require single cell suspensions from fresh tissue isolated directly from surgery, which is often a confined resource and limits the use of clinically well-defined specimens. In addition, duration of surgery is often unknown and maintenance of tissue integrity following tissue resection is not prioritized and further challenges scRNA-seq with a laboratory protocol requiring a minimum of eight hours laboratory work^7,8^. Furthermore, clinical information, treatment response and outcome of the patients are unknown at the time of tissue procurement, and hence expensive and often unnecessary analyses of tumors, not relevant for the study in question, are carried out. In contrast, single-nucleus RNA-sequencing (snRNA-seq) makes it possible to investigate fresh, fixed, or frozen tissue using e.g. the DroNc-seq method, a massively parallel snRNA-seq droplet-based technology, developed by Habib et al.^7^.

snRNA-seq facilitates hypothesis-generating studies and analysis of histologically well-characterized tumor biopsies from patients with long-term clinical follow-up. From a practical point of view, the snRNA-seq method benefits from short processing time compared to scRNA-seq methods and experiments can be carried out when needed. There is a risk of bias when isolating whole tumor cells due to differences in the composition of tumor specimens and the robustness of the cell membrane of tumor cells. This bias is diminished by nuclei isolation because the nuclear membrane is more robust^9^. In addition, it is possible to isolate single nuclei from tissues, where single cell isolation may be difficult^10^. However, we have not found previously published protocols for nuclei isolation^11,12^ applicable for both non-muscle invasive and muscle invasive cancerous bladder tissue, nor adequately gentle for isolation of nuclei at an intact state to maintain nucleus integrity.

Here, we describe the full workflow, from frozen tumor biopsies to snRNA-seq data. The workflow includes a protocol for gentle unbiased nuclei isolation from fresh frozen human bladder tumor biopsies followed by DroNc-seq analysis and optimized library preparation for next generation sequencing. The protocol is robust and highly reproducible across various tumor stages and tumor morphology structures. The nucleus isolation protocol combines optimized tissue dissection using sectioning technique and short incubation time ensuring nuclei isolation from frozen bladder tumor specimens of a quality high enough to be used for snRNA-seq.

## Materials and methods

### Patient samples and processing

Bladder tumor samples were obtained from patients diagnosed with non-muscle invasive bladder cancer (NMIBC) or muscle invasive bladder cancer (MIBC). All patients had provided written informed consent to participate in future research projects before inclusion. The study was approved by The National Committee on Health Research Ethics (#1706291 & #1708266). Tumor samples were obtained from transurethral resection of the bladder or radical cystectomy.

Seven biopsies were processed fresh directly from surgery, 58 samples were embedded in O.C.T., frozen in liquid nitrogen and subsequently stored at −80 °C, and 19 samples were dry frozen in liquid nitrogen without O.C.T. and stored at −80 °C. The tumors represented various tumor stages (31 Ta; 27 T1; 26 T2-4) and tumor grades (26 low grade; 55 high grade; 3 unknown grade).

## Cell culture

Human bladder cancer cell line T24 was obtained from American Type Culture Collection (ATCC-LGC standards, Borås, Sweden) and re-authenticated via STR analysis using the Cell-ID-system (G9500, Promega, Nacka, Sweden). Murine embryonic fibroblast cell line, NIH 3T3, were provided by C. Holmberg, University of Copenhagen, Denmark. Both cell lines were cultivated in Dulbecco’s Modified Eagle Medium (DMEM) (Lonza, cat #BE13-604F) supplemented with 10% Fetal Bovine Serum (FBS) (Gibco, cat #10270-106) and 1% penicillin/streptomycin (Gibco, cat #15140-122) and cultured at 37 °C, 5% CO2 and 90% humidity to a confluence of 70-90%.

### Preparation of Cell Lines for Nuclei Isolation for Nucleus Concentration Analysis and DroNc-seq Experiments

T24 cells and NIH 3T3 cells cultured in separate flasks were washed with 1x PBS (Life Technology, cat #BE17-512F), and treated with 0,05% Trypsin-EDTA (1X) (Gibco, cat #25300-062) for 3-5 min. Trypsin was neutralized with 3 mL of growth medium, and the suspensions were spun down at 500× g for 5 min. at 4 °C. Supernatant was removed, the pellets were washed with 3 mL PBS, and spun down again at 500× g for 5 min. at 4 °C. Following, the supernatant was removed.

#### Nuclei isolation

Below, two protocols for nuclei isolation are described. The first protocol describes isolation of nuclei from cell lines for nucleus concentration analysis. This protocol could not be used for human bladder tumor tissue, since the Nuclei EZ lysis buffer resulted in a high fraction of small and shrunken nuclei (**figure 2G-I**). Using the second protocol, however, decreased the fraction of damaged nuclei and successfully returned isolated round nuclei from tumor tissue (**figure 2A-C**). The second protocol describes isolation of nuclei from T24 human bladder cancer cell line and human muscle invasive and non-muscle invasive bladder tumors prior to DroNc-seq. This latter protocol is more efficient and maintains nucleus integrity (supplementary table 1).

### Nuclei isolation for nucleus concentration analysis

The T24 and NIH 3T3 cell pellets were dissolved in 4 mL ice-cold Nuclei EZ lysis buffer (Sigma Aldrich, cat #NUC-101) and incubated on ice for 5 min. The suspension was centrifuged at 500× g for 5 min. at 4 °C, the supernatant was removed and the pellet resuspended in 4 mL Nuclei Suspension Buffer 1 (NSB1; 1x PBS, 0.01% BSA (New England Biolabs, cat #B9000S) and 0.5 U per µL RNase inhibitor (Nordic Biolabs, cat #30281-1)). The suspension was centrifuged at 500× g for 5 min. at 4 °C, and the supernatant was discarded. The nuclei were resuspended in 2 mL NSB1, and filtered through a 20 μm cell strainer (pluriSelect, cat #43-10020-40). Nuclei suspension was stained 1:1 with Tryphan Blue Solution 0.4% (Roche, cat #5650640001), loaded on a Bürker-Türk counting chamber, and evaluated under a light microscope to ensure properly isolated, intact single nuclei. For the nucleus concentration analysis using DroNc-seq, the nuclei suspensions were diluted in NSB1 to a concentration of 450 nuclei per μL, and the two cell specimens were mixed 1:1 before they were loaded to the DroNc-seq system.

### Nuclei isolation for DroNc-seq and 10x Chromium

Nuclei from a T24 cell line and human bladder tumor biopsies for DroNc-seq experiments were isolated with IgePal lysis buffer (50 mM Tris-HCl pH 8.5 (Sigma Aldrich, cat #T3253-100G), 150 mM NaCl (EMD Millipore, cat #106406), and 1% IGEPAL® CA-630 (Sigma Aldrich, cat #I8896)). Tumor biopsies were sectioned into 50 µm sections on a cryostat. The remaining protocol is identical for T24 cell line and bladder tumor biopsies used for nuclei isolation for DroNc-seq and 10x Chromium. A pellet of T24 cells or sections of tissue were placed in a tube with 5 mL ice cold IgePal lysis buffer and incubated for 3-5 min. The suspension was filtered through a 70 µm cell strainer (Corning, cat #431751) followed by a 50 µm cell strainer (CellTrics, cat #04-004-2327). Nuclei were collected by centrifugation at 500× g for 5 min. at 4 °C. The nuclei were washed in 4 mL Nuclei Suspension Buffer 2 (NSB2; 1x PBS, 0.05% BSA, 0.5 U per µL RNase inhibitor and 1 mM DL-Dithiothreitol solution (Sigma Aldrich, cat #43816)), filtered through a 40 µm cell strainer (Flowmi, cat #15342931) and collected at 500× g for 5 min. at 4 °C. The supernatant was removed and the isolated nuclei was resuspended in an appropriate amount of NSB2. Lyophilized DAPI (Invitrogen, cat #D21490) was dissolved in sterile filtered deionized water to a working solution of 1 µg per mL. Nuclei suspensions was stained 1:1 with either Tryphan Blue Solution or 0.4 % Erythrosin (Sigma Aldrich) in DPBS and DAPI solution, loaded on a Bürker-Türk counting chamber, and evaluated under a light and/or fluorescence microscope to ensure properly isolated, intact singletons. The nuclei were counted and diluted in NSB2 to a final concentration of 453 nuclei per μL for DroNc-seq experiments and 1,000 nuclei per μL for 10x Chromium experiments.

For optimization and comparable experiments of protocols for nuclei isolation, nuclei were also isolated using Nuclei EZ lysis buffer combined with tissue dissectioning by mechanical dissociation with dounce homogenizers (Sigma Aldrich, cat #D8938), semi-automated dissociation with gentleMACS Octo Dissociator with Heaters (Miltenyi Biotec, cat #130-096-427), and a combination of mechanical and semi-automated tissue dissectioning (supplementary table 1, supplementary figure 1).

This method utilizing IgePal lysis buffer for nuclei isolation was the most successful and efficient protocol to yield the highest number of morphologically healthy-looking and visibly intact nuclei isolated (**figure 2A-C**) in relation to minimal use of tissue (**supplementary table 1**), and reduced processing time.

#### Bulk mRNA analysis

T24 cells, tissue sections from a human bladder tumor biopsy, and nuclei, from same batch of both, isolated as described above, were lysed and total RNA purified using RNeasy Mini Kit (Qiagen, cat #74106). mRNA was isolated from 500 ng total RNA using KAPA mRNA Capture Kit (KAPA Biosystems, cat #KK8441), and next generation sequencing libraries were constructed using KAPA mRNA HyperPrep Kit (KAPA Biosystems, cat #KK8544).

Bulk RNA libraries were sequenced on the Illumina NovaSeq 6000 platform using cycle parameters Read 1: 150 bases, Index read 1: 8 bases, Index read 2: 8 bases and Read 2: 150 bases. Salmon^13^ was used to quantify the expression of transcripts using annotation from the Gencode release 33 on genome assembly GRCh38. Transcript-level estimates were imported and summarized at gene-level using the R package tximport v1.20.0 and counts were normalized using the R package edgeR v3.34.1. The R package ggpointdensity v0.1.0 was used to create scatter plots. Each point is colored according to the number of neighboring points. Default settings were used.

### Microfluidic system for DroNc-seq and Drop-seq

To encapsulate nuclei with barcoded beads for DroNc-seq analysis, the Dolomite Bio scRNA-seq system (Dolomite Bio, cat #3200538) with high-speed digital microscope and camera (Dolomite Bio, cat #3200531), and sNuc-seq Chip (85μm Etch Depth, Dolomite Bio, cat #3200607) was used. A loading concentration of 450 nuclei per μL was used for the nucleus concentration analysis, and a loading concentration of 453 nuclei per μL was used for snRNA-seq analysis of bladder tumors on DroNc-seq.

Barcoded beads (Chemgenes, cat #Macosko-2011-10(V+)) was washed, filtered through a 100 µm cell strainer (Corning, cat #431752) followed by a 70 µm cell strainer, and suspended in Drop-seq Lysis Buffer (6% Ficoll PM-400 (GE Healthcare, cat #17-0300-10), 0.2% Sarkosyl (Sigma Aldrich, cat #L7414), 0.02M EDTA (ThermoFisher, cat #AM9261), 0.2 M Tris Ph 7.5 (Invitrogen, cat #15567027), 0.05M Dl-Dithiothreitol solution (Sigma Aldrich, cat #43816)) as described in Habib et al.^7^. The beads were counted using c-chip DHC-F01 counting chambers (NanoEnTek, cat #631-1096), the concentration was corrected to ∼500,000 beads per mL and then they were ready for experiment run. For droplet generation, nuclei and beads were applied for the system at flow rates of 20 μL per min. Droplet Generation Oil for EvaGreen (BioRad QX200™, cat #1864006) was used to encapsulate the droplets at a flow rate of 120 μL per min.

The same microfluidic system was used for Drop-Seq performed on T24 cells for comparison. T24 cells was isolated, trypsinized and washed as described in *Nuclei isolation for DroNc-seq*, and filtered through a 40 μm cell strainer (Corning, cat #431750). The sNuc-seq Chip was replaced by a Droplet Chip 2 (100 μm etch depth, fluorophilic, Dolomite Bio, cat #3200583) and the sample loop was adjusted from 6 m to 10 m in length. Furthermore, a cell concentration of 300,000 cells per mL and a bead concentration of 300,000 beads per mL were loaded for droplet generation at a flow rate of 30 μL per min., and the flow rate for oil was increased to 200 μL per min.

### DroNc-seq and Drop-seq library preparation

Droplet breaking, washes and reverse transcription (RT) was performed^7^. Beads were washed, treated with exonuclease I, resuspended in 1 mL H_2_O, and then counted^7^. Aliquots of 5,000 beads (∼250 STAMPs) were amplified^7^ using the following PCR program: 95 °C for 3 min., next four cycles of 98 °C for 20 s, 65 °C for 45 s, 72 °C for 3 min., then 9 cycles for Drop-seq and 12 cycles for DroNc-seq of 98 °C for 20 s, 67 °C for 20 s, 72 °C for 3 min., and finally 72 °C for 5 min. Supernatants from multiple PCR tubes with the same sample origin were pooled in a 1.5 mL Eppendorf tube, and 0.6× SPRI cleanup (Ampure XP, Beckman Coulter, cat #A63882) was performed. The cleanup procedure was repeated once for DroNc-seq samples. Quality assessment was performed on purified cDNA. Tagmentation and amplification was performed using Nextera XT DNA Library Preparation Kit (Illumina, cat #FC-131-1096) and 600 pg input of each sample. A final 0.6× SPRI cleanup and quality assessment was performed.

### Sequencing of DroNc-seq libraries

Single nuclei libraries were sequenced on the Illumina NovaSeq 6000 platform using a custom Read 1 primer (Read1CustomSeqB see **supplementary table 2**) and cycle parameters Read 1: 26 bases, Index read 1: 8 bases, Index read 2: 0 bases and Read 2: 75 bases. Read 1 reads through 12 bases of nuclei barcode, 8 bases of unique molecular identifier (UMI), 1 base (A/G/C), and then into the poly-T strand. According to recommendations from Illumina, 26 bases in read 1 were sequenced to ensure all Real-Time Analysis calculations are complete.

### Preprocessing of DroNc-seq and Drop-seq data

Processing of the FASTQ files was done as described in the Drop-Seq Core computational Protocol^14^ with a few supplementary steps. First, the reads were sorted and only the ones where the barcodes were correctly formed, i.e. the molecular barcode was followed by a V (a A, C or G) and then the polyT were kept. Then the pipeline was run as described. To summarize: tag the biological read (read2) with the cell barcode and the UMI, filter reads with low quality barcodes, trim 5’ primer sequence, trim 3’ polyA, align the biological read with STAR, add gene/exon and other annotation tags and finally repair substitution errors or indel errors.

For each experiment, we then calculated the expression per barcode for all barcodes with a minimum of 150 genes expressed. The top ranked barcodes (either by the number of genes expressed or by the total read counts associated) were white-listed. The number depended on the theoretical number of single cells expected. We then created different tags where non-white-listed barcodes were changed to a white-listed barcode if the hamming distance between the two barcodes was less or equal to 2 (H2). Finally, for each tag, we recalculated the expression of each gene for all the barcodes in the white-list.

### snRNA-seq using 10x Chromium

For comparison of methods, T24 cells were also subjected to snRNA-seq using the 10x Chromium platform. Nuclei were isolated as described above (*Nuclei Isolation for DroNc-seq and 10x Chromium*), dilated to a concentration of 1,000 nuclei per μL and processed according to the 10x Chromium protocol, Chromium Next GEM Single Cell 3ʹ Reagent Kits v3.1. Libraries were sequenced on an Illumina NovaSeq 6000 platform. Sequencing data were processed with CellRanger from 10x Genomics v7.0.0 using a pre-mrna reference (GRCh38-2020-A) and included introns.

### Comparison of single-cell RNA-seq (Drop-seq) and single-nuclei RNA-seq (DroNc-seq)

We compared the average transcriptomic profiles of single cells (Drop-seq) and single nuclei (DroNc-seq) from a T24 cell line by calculating the average log-transformed UMI counts and the Pearson correlation coefficient. We calculated the differential expression of genes between cells (Drop-seq) and nuclei (DroNc-seq) using the R software package Seurat^15^ v3.1.0. The fraction of mapped reads for single cells (Drop-seq) and single nuclei (DroNc-seq and 10x Chromium) mapping to the exonic region, intergenic region, intronic region and mitochondrial region were also compared for a T24 cell line and three human bladder tumors.

## Results

### Nuclei isolation

A laboratory protocol for snRNA-seq based on nuclei isolation was developed using human bladder tumors comprising both muscle invasive and non-muscle invasive disease (**figure 1**). The cell lysis conditions are paramount for successful isolation of nuclei with an intact nuclear membrane. To establish optimal conditions for this, different lysis buffers (IgePal and Nuclei EZ) and incubation times were tested in combination with different approaches for tissue and cell dissection; sectioning, semi-automated dissociation, manual dissociation with pestles, and semi-automated dissociation combined with manual dissociation with pestles. Following each experiment, the nuclei were counted and carefully examined regarding morphology, nuclei integrity (DNA leakage and blebbing), and isolation efficiency using fluorescence and light microscopy (**supplementary figure 1, supplementary table 1**).

**Figure 1.**
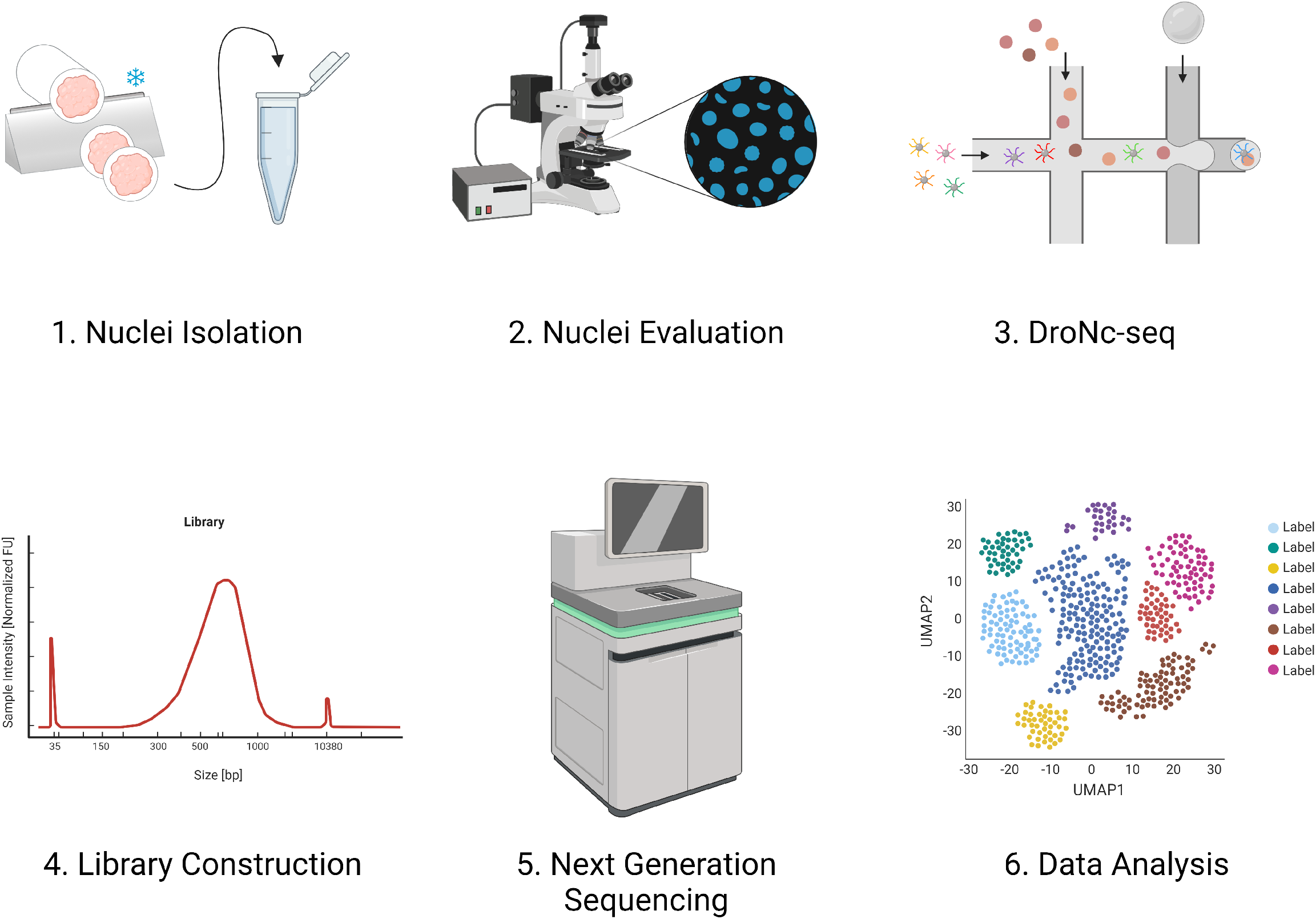
Experimental workflow. 1) Nuclei isolation through tissue dissection by sectioning and incubation in IgePal lysis buffer. 2) Evaluation of DAPI-stained nuclei using fluorescence microscopy. 3) Nuclei encapsulation using DroNc-seq. 4) Construction of libraries. 5) Next generation sequencing of libraries. 6) Data analysis. Created with BioRender.com.

Dissection of the tissue by sectioning was most effective for isolation of a high number of undamaged nuclei relative to the amount of input tumor tissue (mm^3^). This was measured by efficiency (**supplementary table 1, figure 2A-C**). However, manual tissue dissection using pestles was also possible, but time-consuming and released a lot of debris demanding additional washing and filtration steps which can also be critical if a limited amount of tissue is available (**figure 2D-E**). The usage of Nuclei EZ lysis buffer combined with manual dissociation with pestles also damaged many nuclei and increased the amount of debris observed in the microscope (**figure 2F**). The Nuclei EZ lysis buffer was often too harsh on the tissue, leaving no or very few intact nuclei which were shrunken and oblonged (**figure 2G-H**) and had an uneven spikey circumference (**figure 2I**). In contrast, IgePal lysis buffer with the detergent IGEPAL® CA-630 lysed only the plasma membrane leaving intact nuclei which had a smooth surface and a round to oval shape. This is shown for nuclei isolated from human bladder tumors **figure 2A-C** and nuclei isolated from a T24 cell line (**figure 2J-K**). Optimal incubation time in IgePal lysis buffer was 3-5 min. for non-muscle invasive bladder tumors (**figure 2L**) and 5 min. for muscle invasive bladder tumors (**figure 2A-C**). Prolonged incubation for more than 3-5 min. depending on tumor invasiveness, introduced swollen and blebbing nuclei (**figure 2M**) and introduced ruptures of the nuclear membrane resulting in DNA leakage from the nuclei (**figure 2N-P**). Following lysis of the plasma membrane, nuclei were collected by centrifugation and two rounds of wash and filtering were performed to eliminate debris and residual cytoplasmic RNA. Single nuclei suspensions were stained with DAPI and Tryphan Blue Solution or Erythrosin and imaged with fluorescence and light microscopy to ensure an intact nuclear membrane (**figure 2A-C**) and no plasma membrane encapsulated cytoplasmic content. Incomplete isolation and detachment from the plasma membrane were observed when adjusting the incubation time for sectioned tissue with IgePal lysis buffer (**figure 2Q-R**). Furthermore, to ensure having enough nuclei for obtaining 3,000 STAMPs (single nucleus transcriptomes attached to microparticles, i.e. the result of a droplet with the transcriptome of a single nucleus attached to oligos of one bead) we isolated 145,000 nuclei. When approximately 30 mm^3^ tissue or more was used for nuclei isolation, we exceeded this threshold (**supplementary table 1**).

**Figure 2.**
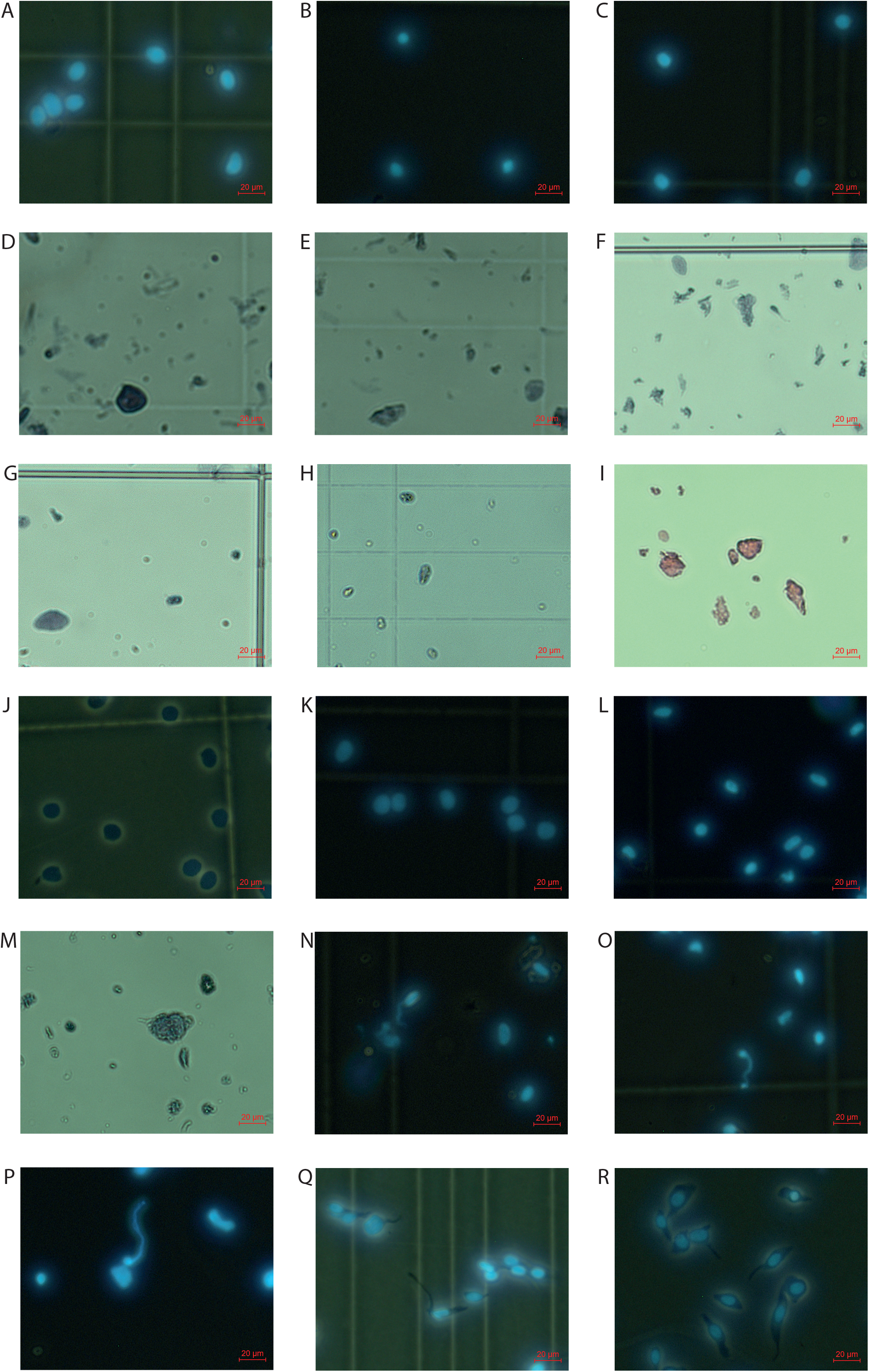
Isolated nuclei. A-C) Undamaged round nuclei isolated from muscle-invasive human bladder tumors. A) Sample 59, B) Sample 64 and C) Sample 71. D-F) High degree of debris observed when nuclei isolation is performed with manual dissociation with pestles combined with D-E) Igepal lysis buffer (sample 34) and F) Nuclei EZ lysis buffer (sample 11). G-I) Nuclei treated with Nuclei EZ lysis buffer were G-H) shrunken and oblonged and had I) a spikey circumference. G) Sample 13, H) Sample 25 and I) Sample 6. J-K) Undamaged round nuclei isolated from a T24 human bladder cancer cell line using 5 min. incubation in IgePal Lysis Buffer. L) Undamaged round nuclei isolated from a non-muscle invasive human bladder tumor (sample 76). M) Swollen and blebbing nuclei resulting from prolonged incubation of a human bladder tumor (sample 27). N-P) DNA leakage from prolonged incubation of human bladder tumors for N-O) > 5 min. and P) > 3 min. N) Sample 43, O) Sample 50 and P) Sample 70. Q-R) Incomplete nucleus isolation from human bladder tumors with remains of the cytoplasmic membrane. Q) Sample 56 and R) Sample 57. Nuclei are stained 1:1 with DAPI, Tryphan Blue Solution or Erythrosin and evaluated through a fluorescence or light microscope.

In summary, at least 30 mm^3^ tissue was used for nuclei isolation by sectioning on a cryostat. This was followed by incubation in IgePal lysis buffer for 3-5 min. to disrupt the outer cytoplasmic membrane and isolate the highest number of whole intact nuclei (>145,000 nuclei) regardless of tumor stage and tissue structure.

### Bulk nuclei and bulk whole cell gene expression comparison

Prior to single nucleus analysis, we inspected whether the protocol for nuclei isolation changed the transcriptomic profile beyond the compartment restricted RNA species. We performed bulk mRNA-sequencing of the nuclear and cellular fractions from a bladder tumor and from a T24 human bladder cancer cell line. The nuclear fractions were isolated as described above and evaluated under a fluorescence microscope using DAPI to ensure properly isolated nuclei without remains of the cytoplasmic content. Nuclei, cells and tissue were lysed and bulk mRNA libraries were generated and sequenced. The bulk mRNA-sequencing data showed a relatively high correlation between gene expression from the nuclear and cellular fractions from both samples (**Pearson R > 0.6, p < 2.2e-16, figure 3**). Mitochondrial genes were higher expressed in the cellular fraction for T24 compared to the nuclear fraction (**figure 3A**), as seen in previous studies^16^. These differences in mitochondrial gene expression were not identified for the tumor biopsy analysis, which could be caused by an overall higher cell cycle and energy requirement in pure cancer cells from the T24 cell line. Nuclear noncoding RNAs like *MALAT1, NEAT1*, and *XIST* were found to be highly expressed in both bulk nuclei mRNA and bulk cell mRNA (figure 3A-B), probably because the nuclei noncoding RNAs are present in both fractions analyzed. These data suggest that the nuclei isolation protocol introduces limited bias to the transcriptome analysis.

**Figure 3.**
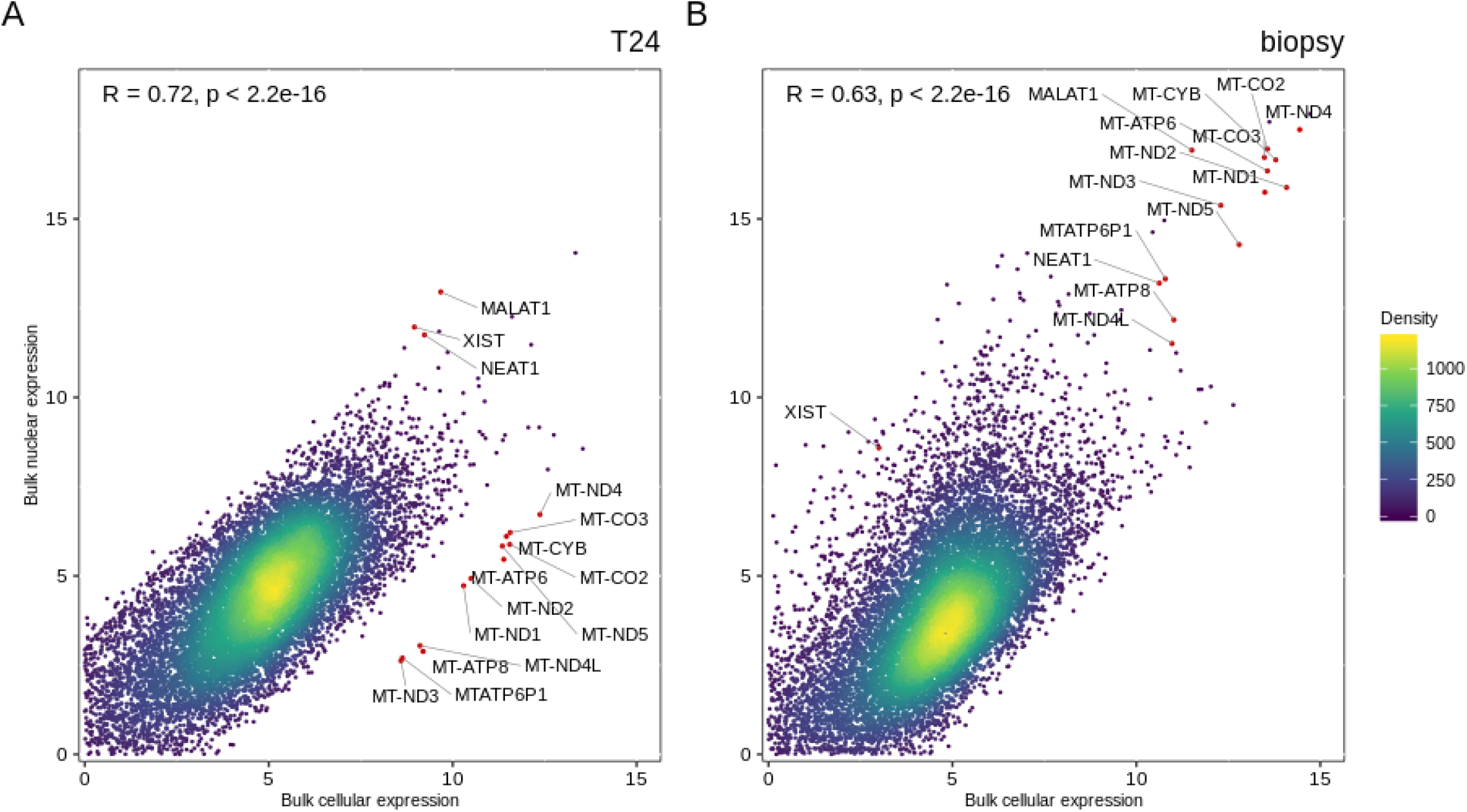
Concordance between gene expression profiles for nuclei and cells. Scatterplot of gene expression [log(counts+1)] from fractions of cellular and nuclear bulk mRNA from A) T24 human bladder cancer cell line **R = 0.72, p < 2.2e-16** and B) human muscle-invasive bladder tumor **R = 0.63, p < 2.2e-16**. Pearson correlation was used to determine the correlation coefficient R and p-value. Density indicates neighboring data points. Red dots indicate deviating genes highly expressed in one fraction but not the other.

### Nucleus concentration analysis

Studies on Drop-seq and DroNc-seq dilute cells and nuclei to poisson-limiting concentrations in droplets (nuclei in 5% of droplets)^7,8^. The manufacturer for our system recommended a nucleus concentration of 453 nuclei per μL. However, the optimal concentration may differ depending on the nucleus size, hence a nucleus concentration analysis was carried out to determine the appropriate nucleus concentration to avoid too many doublets, i.e. droplets that encapsulate two nuclei resulting in a mixture of their RNA and thereby a false RNA profile. To estimate this, a mixing experiment was carried out using nuclei isolated from two different cultured cell lines (T24 human bladder cancer cell line and NIH 3T3 murine cell line) mixed in ratio 1:1. Based on earlier reports^7^, an experiment with 450 nuclei per μL was performed. The identity of a particular nucleus was defined as human or murine origin if more than 95% of the transcripts harboring the same cell barcode mapped to the human or murine reference genome, respectively. A droplet was considered to have a mixed origin (i.e. a droplet containing one human nuclei combined with one murine nuclei) if less than 95% of the transcripts mapped to one of the reference genomes. The experiment revealed a doublet rate of 4.09% when using a concentration of 450 nuclei per μL (**figure 4**). A concentration of approximately 453 nuclei per μL was used for DroNc-seq analysis of human bladder tumors.

**Figure 4.**
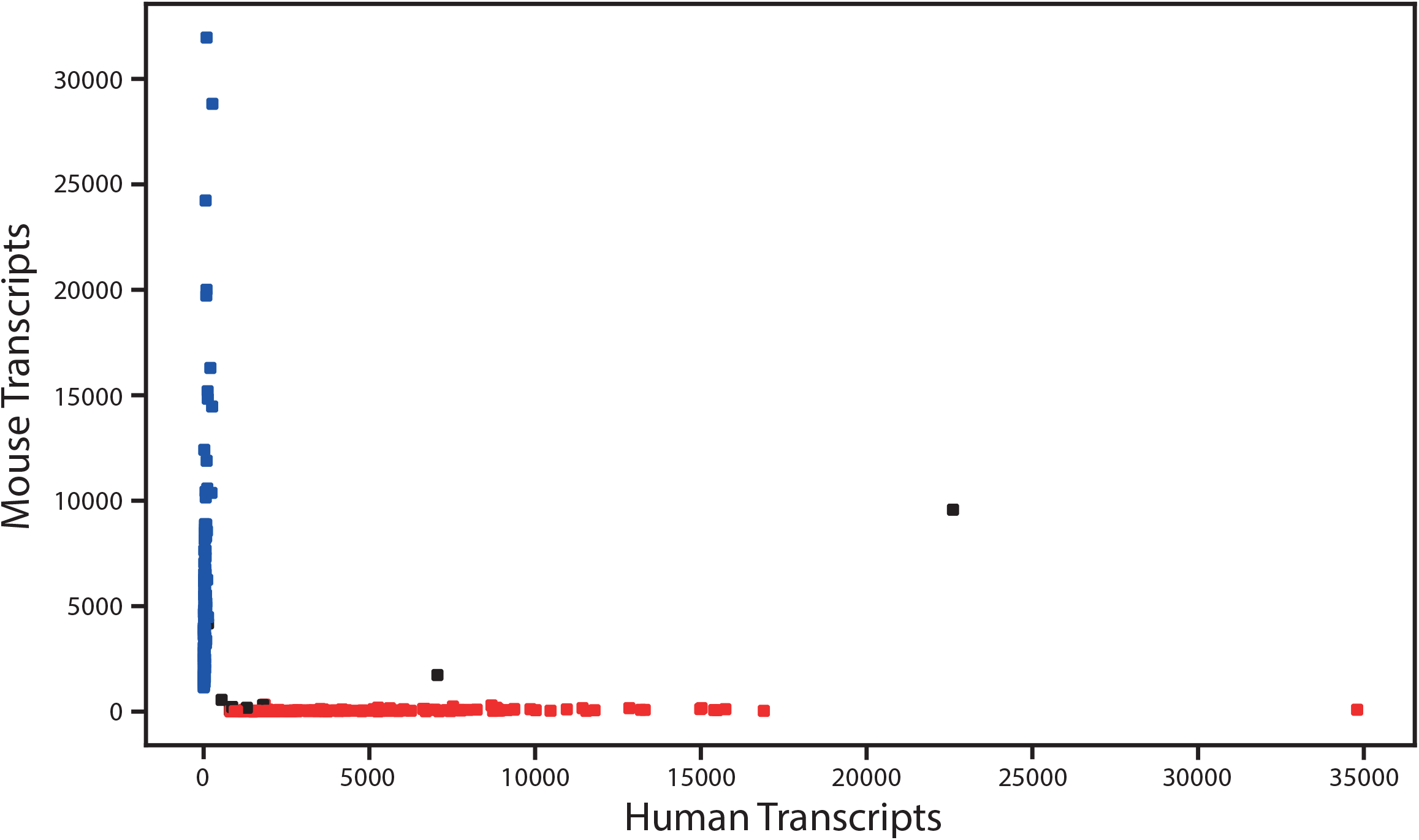
Nucleus concentration analysis. Nucleus identity of the mixing experiment using a nucleus concentration of 450 nuclei per μL. 204 unique murine nuclei, 179 unique human nuclei, and 8 mixed nuclei were identified. A droplet was considered as a doublet (mixed origin) if <95% of the transcripts mapped to one of the reference genomes. Blue dots: droplets with murine nuclei. Red dots: droplets with human nuclei. Black dots: droplets with human and mouse nuclei. Cut-off values: min. 1 read per gene, min. 200 genes per nuclei, >95% reads must map to the human or murine reference genome, respectively, to define a cell as unique.

### Gene expression concordance analysis for single nuclei and single cells

To investigate whether the transcriptional profiles of single nuclei were representative of whole cells, scRNA and snRNA-seq were applied on isolated T24 cells and nuclei, respectively. ScRNA-seq and snRNA-seq data was processed using the Macosko pipeline^14^ with the addition of a few supplementary steps described in the method section under: *Preprocessing of DroNc-seq and Drop-seq data*. We compared the average expression of each gene (log-transformed counts) obtained for single nuclei and single cells to investigate if the nuclear transcriptomic profile contained a high degree of similarity to the cellular expression profile. We found a correlation of **R = 0.59, p < 2.2e-16** between the average single cell and single nucleus transcriptome (**figure 5A**). Genes significantly higher expressed in nuclei (*NEAT1, MALAT1* and lncRNAs, *PAX8-AS1* and *XIST*) or cells (*MT-RNR2, RPS3, RPL8, RPL11*, and *RPL19*) were consistent with the compartment restricted RNA species.

**Figure 5.**
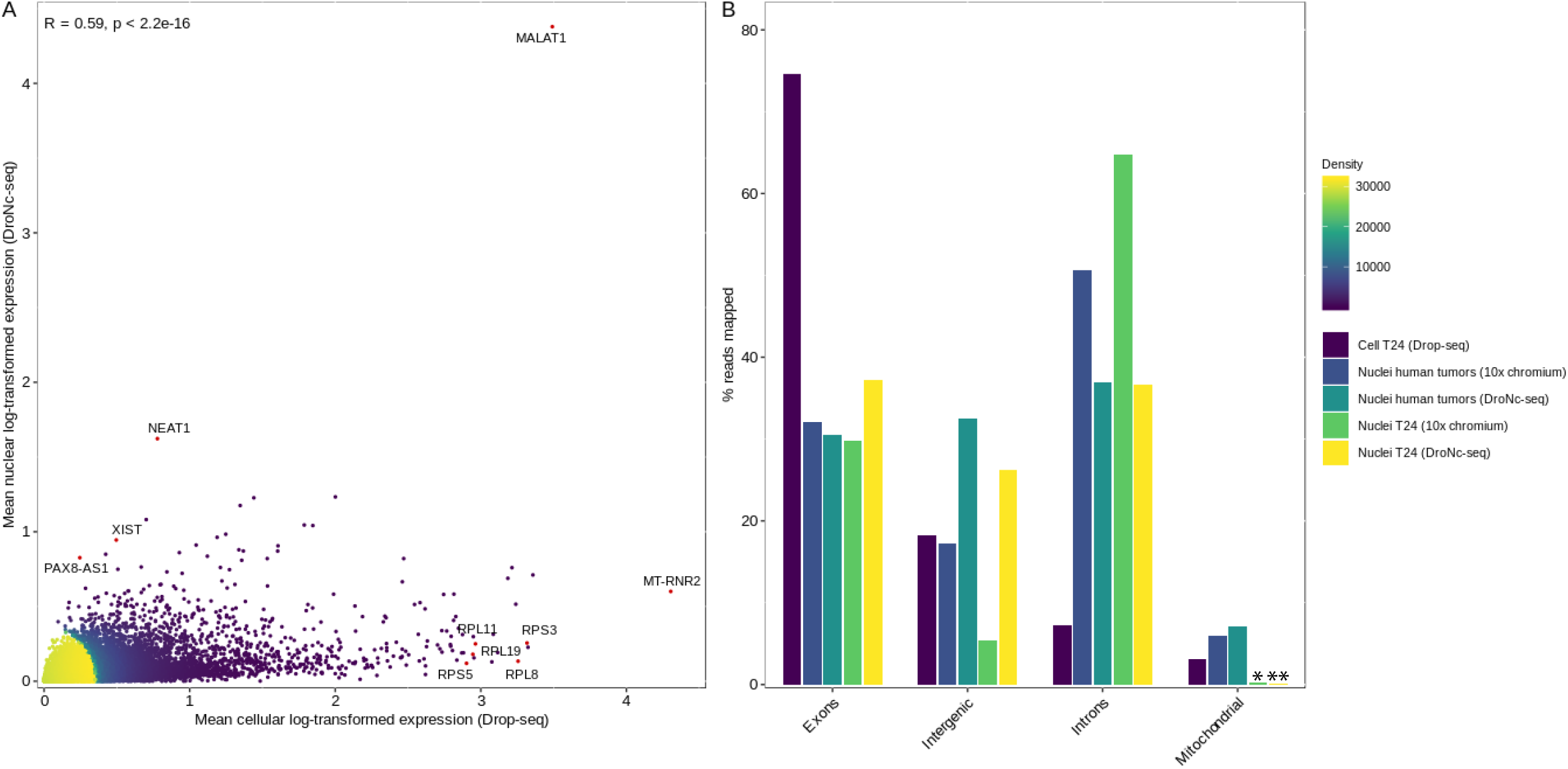
Comparison of gene expression in single nuclei and single cells. A) Scatterplot comparing average gene expression (log-transformed) for single nucleus and single cell data from T24 human bladder cancer cell line. **R = 0.59, p < 2.2e-16**. Pearson correlation was used to determine the correlation coefficient R and p-value. Selected genes significantly expressed in either cells or nuclei are marked in red. Density indicates neighboring data points. B) Fraction of reads mapping to exonic, intronic, intergenic and mitochondrial regions (out of the reads mapped to the genome) for T24 cells (Drop-seq), T24 nuclei and human tumors (10x Chromium and DroNc-seq). % reads for human tumors is an average of sample 48, 72, and 84. Mitochondrial expression in * T24 nuclei (10x Chromium) was 0.21 % and in ** T24 nuclei (DroNc-seq) was 0.11 %.

We also compared the fraction of reads mapping to exonic, intronic and mitochondrial regions for scRNA-seq (Drop-seq) and snRNA-seq data obtained from two different platforms, DroNc-seq and 10x Chromium. As expected; for single cells (Drop-seq) a large fraction of reads (74.57%) mapped to exons, whereas for single nuclei the corresponding fractions of reads mapping to exons was lower (30.57-37.19% for DroNc-seq and 29.84-32.08% for 10x Chromium) (**figure 5B**). Nonetheless, a low fraction of reads mapped to introns (7.18%) for single cells (Drop-seq) compared to single nuclei (36.58-36.93% for DroNc-seq and 50.69-64.76% for 10x Chromium) (**figure 5B**). The observed fractions of intron- and exon-mapped reads in the nuclei may reflect the maturation process of RNA in the nuclei^12^. Furthermore, we show that nuclei isolated with this optimized protocol can be applied on different platforms for single nuclei analysis (**figure 5B**). The fraction of exons for nuclei were comparable between the two platforms but 10x Chromium seemed to capture a higher percentage of introns compared to DroNc-seq. Oligos on beads from Chemgenes and 10x gel beads are both constructed with a 30-dNTP to capture the mRNA, which therefore does not account for the difference. A possible explanation could be differences between how the bioinformatic pipelines assign reads to exons versus introns.

## Discussion

Here we describe a protocol for snRNA-seq including a robust procedure for gentle nuclei isolation from fresh frozen tissue using dissection of tissue by sectioning and incubation in IgePal lysis buffer. Comparison of gene expression profiles from nuclei and cells showed a high concordance and genes known to be specifically expressed in different cellular compartments were identified^16,17^. The fraction of reads obtained from either nuclei or cells were enriched according to the origin of the RNA analyzed, which is congruent with previous studies^7,12^.

This protocol for nucleus isolation has been developed and optimized for human bladder cancer tissue. If the protocol is used to isolate nuclei from other types of tumor tissue or species, optimization is recommended in order to find the optimal incubation time, as we observed a difference between tissues from NMIBC and MIBC.

The majority of previous studies within the field have investigated single cells from fresh tissue, which may cause a challenge in laboratory workflow and lack of information on the samples being analyzed - especially in clinical studies. It is not possible to analyze single cells isolated from frozen biobanked material since the RNA integrity in biopsies decreases drastically if the sample is thawed after it has been snapfrozen because of RNase activity^18^. This stresses the advantage that snRNA-seq has over scRNA-seq because already biobanked material and tissue difficult to dissociate into single cells can be analyzed. Explorative studies can utilize this and be designed for patients with well described clinical and histopathological information and known clinical outcomes.

A previous study comparing scRNA-seq and snRNA-seq showed that fewer immune cells were recovered from frozen mouse kidney tissue compared to fresh tissue (0.73% and 6.03%, respectively) with an underrepresentation of T-, B-, and natural killer-lymphocytes in their single nuclei study^19^. However, in another study, Gouin et al. performed snRNA-seq on fresh frozen bladder tumor samples and identified a fraction of immune cells (5%) comprising T-cells, dendritic cells, macrophages, and B-cells defined by classic immune marker genes^4^. A third study on human cervical squamous cell carcinoma also performing snRNA-seq identified a diverse population of immune cells as well^20^. Although they identify immune cells, these studies may underestimate the true fraction of immune cells present in their natural conditions after all. It is unknown whether the variation in immune populations identified is tissue specific, caused by heterogeneity within the tissue or caused by preservation methods. In comparison to DroNc-seq, 10x genomics has also developed a droplet-based platform (Chromium) for single cell and single nucleus analysis. A comparative study of scRNA-seq and snRNA-seq methods showed that 10x Chromium has a higher performance with a notable increase in data per nuclei compared to the DroNc-seq system^10^. This could be due to better error suppression in nuclei barcodes and/or the utilization of elastic beads. Furthermore, the capture rate of 10x Chromium is 60% whereas the capture rate of DroNc-seq is 5%^7^, putting a higher demand on the amount of input material needed. However, while the 10x Chromium platform is less time-consuming and a more standardized system, the DroNc-seq system is more cost-effective, which is favorable for high-throughput studies that include multiple samples.

In conclusion, we successfully optimized a robust and reproducible protocol for nuclei isolation applicable across various tumor stages and tumor morphology structures. This nuclei isolation protocol combined with our optimized library preparation for next generation sequencing provides the full workflow for snRNA-seq of human bladder tumors, from frozen tumor biopsies to data ready for analysis.

## Acknowledgements

We would like to thank all technical personnel at Department of Molecular Medicine and Department of Urology and Oncology, Aarhus University Hospital, for sample handling and processing. We would like to thank GenomeDK and Aarhus University for providing computational resources that contributed to these research results.

## Competing interests

Lars Dyrskjøt has sponsored research agreements with C2i, AstraZeneca, Natera, Photocure, and Ferring; has an advisory/consulting role at Ferring and UroGen; and is Chairman of the Board in BioXpedia A/S.

Jørgen Bjerggaard Jensen is proctor for Intuitive Surgery; is a member of advisory board for Olympus Europe, Ambu, Cepheid, and Ferring; and has sponsored research agreements with Medac, Photocure ASA, Cepheid, and Ferring.

## Data availability

The raw sequencing data generated in this study are not publicly available as this compromise patient consent and ethics regulations in Denmark. Processed non-sensitive data are available upon reasonable request from the corresponding author.

## Author contributions

Conceptualization S.S.S., I.N., L.D.; Data Curation S.S.S., S.V.L., P.L., M.K.; Formal Analysis S.S.S., S.V.L., P.L.; Funding Acquisition L.D.; Investigation S.S.S., I.N.; Methodology S.S.S., I.N., L.D.; Project Administration L.D.; Resources J.B.J., L.D.; Supervision L.D.; Visualization S.S.S., M.K.; Writing – Original Draft: S.S.S., I.N., S.V.L., L.D.; Writing – Review & Editing: all authors.

## Funding information

Independent Research Fund Denmark, The Novo Nordisk Foundation, Aarhus University (AUFF NOVA), The Leo & Anne Albert Institute for Bladder Cancer Care and Research.

## Figure legends

**Supplementary Figure 1 Experimental workflows tested for optimization of nuclei isolation**. Three types of tissue conditions (n = 84), fresh, fresh frozen and dry frozen, was tested in combination with various approaches for tissue dissection by sectioning, manual dissection (pestles), semi-automated dissection (gentleMACS), and semi-automated dissection (gentleMACS) combined with manual dissection (pestles). These combinations were further combined with either IgePal lysis buffer or Nuclei EZ lysis buffer. Beneath each workflow, it is stated how many experiments have been performed and how large the success rate was (experiments obtaining >145,000 nuclei). Created with BioRender.com.

**Supplementary Table 1 Experimental overview of workflows tested during optimization of nuclei isolation and results**. Nucleus integrity is evaluated by visible leakage of nuclear DNA or blebbing (yes: nucleus integrity intact, no: visible DNA leakage or blebbing). Gray: not enough nuclei isolated for downstream single-nucleus RNA-sequencing analysis. N/A: no nuclei detected with light- or fluorescence microscopy.

**Supplementary Table 2 Primers used throughout DroNc-seq and Drop-seq experiments**. V: A, C or G. r indicates a ribose-phosphate backbone. * indicate a phosphorothioate bond between two bases. TSO: template switch oligo.

## Notes

### Summary of Updates

The manuscript has been thoroughly revised.

