## Supplementary Figure and Tables for "Improved Protocol for Single-Nucleus RNA-sequencing of Frozen Human Bladder Tumor Biopsies"

### Supplementary figure 1

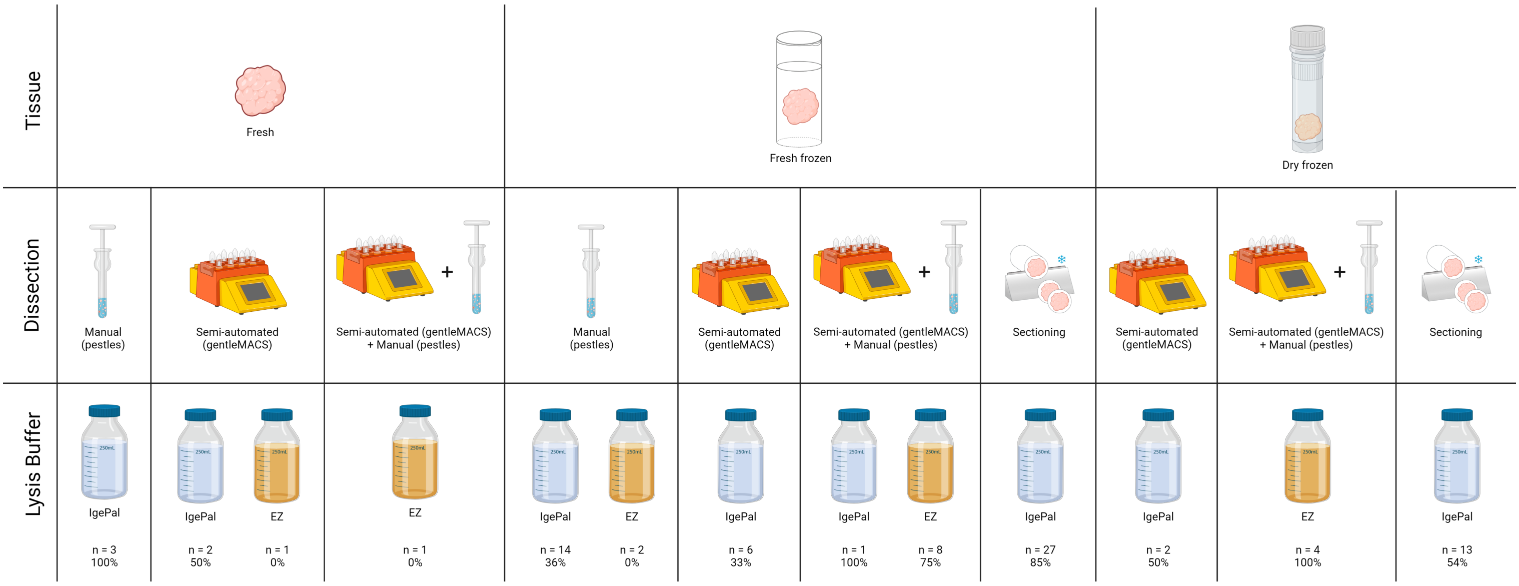

Supplementary table 1

| Sample | Stage | Grade | Type | Tissue volume ($mm^3$) | Lysis buffer | Dissection | Incubation (min.) | Total cells | Nucleus morphology | Nucleus integrity intact | Efficiency (nuclei/$mm^3$ tissue) |
|---|---|---|---|---|---|---|---|---|---|---|---|
| 1 | T2-4 | high grade | fresh frozen | 336.0 | EZ Lysis Buffer | manual (pestles) | 10 | 0 | N/A | N/A | 0 |
| 2 | T2-4 | high grade | dry frozen | 440.0 | EZ Lysis Buffer | semi-automated (gentleMACS) + manual (pestles) | 10 | 10916000* | round, oblong, blebbing | no | 24809 |
| 3 | T1 | high grade | fresh frozen | 126.0 | EZ Lysis Buffer | semi-automated (gentleMACS) + manual (pestles) | 10 | 642000* | round, spikey surface, shrunken | no | 5095 |
| 4 | T1 | high grade | dry frozen | 128.0 | EZ Lysis Buffer | semi-automated (gentleMACS) + manual (pestles) | 10 | 3212000* | round, spikey surface | no | 25094 |
| 5 | T1 | high grade | dry frozen | 72.0 | EZ Lysis Buffer | semi-automated (gentleMACS) + manual (pestles) | 10 | 1500000* | round, spikey surface | no | 20833 |
| 6 | Ta | low grade | dry frozen | 22.5 | EZ Lysis Buffer | semi-automated (gentleMACS) + manual (pestles) | 10 | 1390000* | oblong, spikey surface | no | 61778 |
| 7 | Ta | low grade | fresh frozen | 12.0 | EZ Lysis Buffer | semi-automated (gentleMACS) + manual (pestles) | 10 | 290000* | round, blebbing | no | 24167 |
| 8 | Ta | low grade | fresh frozen | 36.0 | EZ Lysis Buffer | semi-automated (gentleMACS) + manual (pestles) | 10 | 675000 | round, oblong | no | 18750 |
| 9 | Ta | low grade | fresh frozen | 30.0 | EZ Lysis Buffer | semi-automated (gentleMACS) + manual (pestles) | 10 | 1186600 | N/A | N/A | 39553 |
| 10 | Ta | high grade | fresh frozen | 40.0 | EZ Lysis Buffer | semi-automated (gentleMACS) + manual (pestles) | 10 | 0 | N/A | N/A | 0 |
| 11 | Ta | high grade | fresh frozen | 80.0 | EZ Lysis Buffer | semi-automated (gentleMACS) + manual (pestles) | 10 | 550000 | round, oblong | yes | 6875 |
| 12 | Ta | high grade | fresh frozen | 6.0 | EZ Lysis Buffer | semi-automated (gentleMACS) + manual (pestles) | 5 | 0 | round | yes | 0 |
| 13 | T1 | N/A | fresh | 189.0 | EZ Lysis Buffer | semi-automated (gentleMACS) + manual (pestles) | 10 | 40000 | round, oblong, shrunken | yes | 212 |
| 14 | Ta | high grade | fresh frozen | 20.0 | EZ Lysis Buffer | semi-automated (gentleMACS) + manual (pestles) | 5 | 2824000 | round, oblong | no | 141200 |
| 15 | Ta | low grade | fresh | 528.0 | EZ Lysis Buffer | semi-automated (gentleMACS) | 5 | 0 | round | yes | 0 |
| 16 | T1 | high grade | fresh frozen | 12.0 | IgePal Lysis Buffer | semi-automated (gentleMACS) | 10 | 17200* | round | no | 1433 |
| 17 | T1 | high grade | fresh frozen | 12.0 | IgePal Lysis Buffer | manual (pestles) | 30 | 0 | N/A | N/A | 0 |
| 18 | T2-4 | high grade | fresh frozen | 96.0 | IgePal Lysis Buffer | manual (pestles) | 30 | 303400* | round | no | 3160 |
| 19 | Ta | low grade | fresh frozen | 8.0 | IgePal Lysis Buffer | manual (pestles) | 30 | 0 | round | yes | 0 |
| 20 | T1 | high grade | fresh frozen | 0.1 | IgePal Lysis Buffer | manual (pestles) | 30 | 19700* | round, shrunken | yes | 197000 |
| 21 | Ta | high grade | fresh frozen | 2.5 | IgePal Lysis Buffer | manual (pestles) | 30 | 0 | N/A | N/A | 0 |
| 22 | Ta | low grade | fresh frozen | 2.0 | IgePal Lysis Buffer | manual (pestles) | 30 | 0 | N/A | N/A | 0 |
| 23 | T2-4 | high grade | fresh frozen | 105.0 | IgePal Lysis Buffer | manual (pestles) | 30 | 721000* | round | no | 6867 |
| 24 | T2-4 | high grade | fresh frozen | 108.0 | IgePal Lysis Buffer | manual (pestles) | 30 | 710000* | round | yes | 6574 |
| 25 | T2-4 | high grade | fresh frozen | 168.0 | EZ Lysis Buffer | manual (pestles) | 5 | 16500* | shrunken | no | 98 |
| 26 | Ta | low grade | fresh frozen | 25.0 | IgePal Lysis Buffer | semi-automated (gentleMACS) | 2 | 67800* | round | yes | 2712 |
| 27 | T2-4 | high grade | fresh frozen | 280.0 | IgePal Lysis Buffer | manual (pestles) | 10 | 303000* | round, oblong, shrunken, blebbing | no | 1082 |
| 28 | T2-4 | high grade | fresh frozen | 351.0 | IgePal Lysis Buffer | semi-automated (gentleMACS) | 10 | 139000* | round, oblong, shrunken | no | 396 |
| 29 | T2-4 | high grade | fresh frozen | 16.0 | IgePal Lysis Buffer | semi-automated (gentleMACS) + manual (pestles) | 30 | 168000 | round, oblong | no | 10500 |
| 30 | T2-4 | N/A | fresh frozen | 6.0 | IgePal Lysis Buffer | semi-automated (gentleMACS) | 30 | 0 | N/A | N/A | 0 |
| 31 | T2-4 | high grade | fresh | 5000.0 | IgePal Lysis Buffer | semi-automated (gentleMACS) | 30 | 0 | N/A | N/A | 0 |
| 32 | T2-4 | high grade | fresh frozen | 9.0 | IgePal Lysis Buffer | manual (pestles) | 30 | 129000 | round, oblong, shrunken | no | 14333 |
| 33 | T1 | high grade | fresh frozen | 36.0 | IgePal Lysis Buffer | manual (pestles) | 30 | 62700 | round, oblong, shrunken | no | 1742 |
| 34 | T1 | N/A | fresh frozen | 189.0 | IgePal Lysis Buffer | manual (pestles) | 30 | 766500 | round, oblong, shrunken | no | 4056 |
| 35 | T1 | high grade | fresh frozen | 40.0 | IgePal Lysis Buffer | manual (pestles) | 30 | 0 | N/A | N/A | 0 |
| 36 | T2-4 | high grade | fresh frozen | 30.0 | IgePal Lysis Buffer | manual (pestles) | 30 | 55500 | oblong | no | 1850 |
| 37 | T1 | high grade | fresh | 252.0 | IgePal Lysis Buffer | manual (pestles) | 30 | 479250 | round, oblong, shrunken | no | 1902 |
| 38 | Ta | low grade | fresh | 600.0 | IgePal Lysis Buffer | manual (pestles) | 30 | 7234000* | oblong, shrunken | no | 12057 |
| 39 | Ta | low grade | fresh | 600.0 | IgePal Lysis Buffer | manual (pestles) | 20 | 7078000* | round, oblong, shrunken | no | 11797 |
| 40 | Ta | low grade | fresh | 600.0 | IgePal Lysis Buffer | semi-automated (gentleMACS) | 10 | 6133000* | round, oblong | yes | 10222 |
| 41 | T1 | high grade | fresh frozen | 24.0 | IgePal Lysis Buffer | sectioning | 10 | 140000 | round | no | 5833 |
| 42 | T1 | high grade | fresh frozen | 24.0 | IgePal Lysis Buffer | sectioning | 10 | 119000 | round | no | 4958 |
| 43 | T1 | high grade | fresh frozen | 162.0 | IgePal Lysis Buffer | semi-automated (gentleMACS) | 10 | 528000 | round, oblong | no | 3259 |
| 44 | T1 | high grade | fresh frozen | 140.0 | IgePal Lysis Buffer | sectioning | 5 | 1035000 | round, oblong | yes | 7393 |
| 45 | T2-4 | high grade | fresh frozen | 40.0 | IgePal Lysis Buffer | sectioning | 10 | 355000 | oblong, shrinking | no | 8875 |
| 46 | T2-4 | high grade | fresh frozen | 80.0 | IgePal Lysis Buffer | sectioning | 10 | 334640 | round, oblong, shrinking | no | 4183 |
| 47 | T2-4 | high grade | fresh frozen | 330.0 | IgePal Lysis Buffer | semi-automated (gentleMACS) | 10 | 1355000 | round, oblong | no | 4106 |
| 48 | T2-4 | high grade | fresh frozen | 140.0 | IgePal Lysis Buffer | sectioning | 5 | 1347500 | round | no | 9625 |
| 49 | Ta | low grade | dry frozen | 12.8 | IgePal Lysis Buffer | sectioning | 10 | 76000 | round | yes | 5938 |
| 50 | Ta | low grade | dry frozen | 24.0 | IgePal Lysis Buffer | semi-automated (gentleMACS) | 10 | 92500 | round | no | 3854 |
| 51 | T2-4 | high grade | dry frozen | 19.2 | IgePal Lysis Buffer | sectioning | 10 | 120000 | round | no | 6250 |
| 52 | T2-4 | high grade | dry frozen | 105.0 | IgePal Lysis Buffer | semi-automated (gentleMACS) | 10 | 301000 | round | no | 2867 |
| 53 | T2-4 | high grade | dry frozen | 4.0 | IgePal Lysis Buffer | sectioning | 5 | 0 | N/A | N/A | 0 |
| 54 | Ta | low grade | dry frozen | 4.8 | IgePal Lysis Buffer | sectioning | 5 | 0 | N/A | N/A | 0 |
| 55 | Ta | low grade | dry frozen | 5.0 | IgePal Lysis Buffer | sectioning | 5 | 0 | N/A | N/A | 0 |
| 56 | Ta | low grade | dry frozen | 30.0 | IgePal Lysis Buffer | sectioning | 5 | 90500 | round | yes | 3017 |
| 57 | Ta | low grade | dry frozen | 36.0 | IgePal Lysis Buffer | sectioning | 5 | 2782500 | round | yes | 77292 |
| 58 | T1 | high grade | dry frozen | 30.0 | IgePal Lysis Buffer | sectioning | 5 | 1532000 | round | yes | 51067 |
| 59 | T1 | high grade | dry frozen | 65.0 | IgePal Lysis Buffer | sectioning | 5 | 982300 | round | yes | 15112 |
| 60 | T2-4 | high grade | fresh frozen | 22.0 | IgePal Lysis Buffer | sectioning | 5 | 249000 | round | yes | 11318 |
| 61 | T2-4 | high grade | fresh frozen | 36.0 | IgePal Lysis Buffer | sectioning | 5 | 14500 | round | yes | 403 |
| 62 | T2-4 | high grade | fresh frozen | 27.2 | IgePal Lysis Buffer | sectioning | 5 | 59750 | round | yes | 2197 |
| 63 | T2-4 | high grade | fresh frozen | 77.0 | IgePal Lysis Buffer | sectioning | 5 | 2432000 | round, oblong | no | 31584 |
| 64 | T2-4 | high grade | fresh frozen | 88.2 | IgePal Lysis Buffer | sectioning | 5 | 262500 | round | yes | 2976 |
| 65 | Ta | low grade | dry frozen | 45.0 | IgePal Lysis Buffer | sectioning | 5 | 1984375 | round, oblong | no | 44097 |
| 66 | T1 | high grade | fresh frozen | 45.0 | IgePal Lysis Buffer | sectioning | 5 | 789062 | round | yes | 17535 |
| 67 | T1 | high grade | fresh frozen | 45.0 | IgePal Lysis Buffer | sectioning | 5 | 218750 | round | yes | 4861 |
| 68 | Ta | low grade | dry frozen | 22.5 | IgePal Lysis Buffer | sectioning | 5 | 284375 | round | yes | 12639 |
| 69 | Ta | low grade | fresh frozen | 18.0 | IgePal Lysis Buffer | sectioning | 5 | 284375 | round | yes | 15799 |
| 70 | T1 | high grade | fresh frozen | 45.0 | IgePal Lysis Buffer | sectioning | 5 | 398437 | round, oblong | no | 8854 |
| 71 | T1 | high grade | fresh frozen | 45.0 | IgePal Lysis Buffer | sectioning | 5 | 1639102 | round | yes | 36424 |
| 72 | T1 | low grade | dry frozen | 50.0 | IgePal Lysis Buffer | sectioning | 5 | 2108516 | round | yes | 42170 |
| 73 | T1 | low grade | dry frozen | 44.0 | IgePal Lysis Buffer | sectioning | 5 | 324219 | round | yes | 7369 |
| 74 | T1 | high grade | fresh frozen | 45.6 | IgePal Lysis Buffer | sectioning | 5 | 962500 | round | yes | 21107 |
| 75 | T1 | high grade | fresh frozen | 43.8 | IgePal Lysis Buffer | sectioning | 3 | 1062500 | round, oblong | no | 24286 |
| 76 | Ta | low grade | fresh frozen | 45.0 | IgePal Lysis Buffer | sectioning | 3 | 4878828 | round, oblong | yes | 108418 |
| 77 | Ta | low grade | fresh frozen | 48.0 | IgePal Lysis Buffer | sectioning | 3 | 1185078 | round, oblong | yes | 24689 |
| 78 | T1 | high grade | fresh frozen | 45.0 | IgePal Lysis Buffer | sectioning | 3 | 351000 | round, oblong | no | 7800 |
| 79 | Ta | high grade | fresh frozen | 44.8 | IgePal Lysis Buffer | sectioning | 3 | 1931523 | round, oblong | yes | 43114 |
| 80 | T1 | high grade | fresh frozen | 37.5 | IgePal Lysis Buffer | sectioning | 3 | 433594 | round | yes | 11563 |
| 81 | Ta | high grade | fresh frozen | 20.0 | IgePal Lysis Buffer | sectioning | 3 | 1097656 | round | yes | 54883 |
| 82 | Ta | low grade | fresh frozen | 30.0 | IgePal Lysis Buffer | sectioning | 3 | 2490234 | round | yes | 83008 |
| 83 | Ta | low grade | fresh frozen | 25.0 | IgePal Lysis Buffer | sectioning | 3 | 1151563 | round | yes | 46063 |
| 84 | T2-4 | high grade | fresh frozen | 45.0 | IgePal Lysis Buffer | sectioning | 5 | 1167188 | round | yes | 25938 |

* Nuclei were counted on Cedex. Uncertainty of ± 0.067-2.528.

Supplementary table 2

| **Name/Oligo** | **Sequence** | **Conc.** | **Source** |
|---|---|---|---|
| Macosko-2011-10(V+) beads | 5’- Toyopearl-linker-beads-TTTTTTTAAG CAG TGG TAT CAA CGC AGA GTAC JJJJJJJJJJJJ NNNNNNNN V TTTTTTTTTTTTTTTTTTTTTTTTTTTTTT-3’ | 300,000 beads/µl | ChemGenes |
| TSO | AAG CAG TGG TAT CAA CGC AGA GTG AAT rGrGrG | 50 µM | IDT |
| SMART PCR Primer | AAG CAG TGG TAT CAA CGC AGA GT | 100 µM | IDT |
| New-P5-SMART PCR hybrid oligo | AAT GAT ACG GCG ACC ACC GAG ATC TAC ACG CCT GTC CGC GGA AGC AGT GGT ATC AAC GCA GAG T*A*C | 10 µM | IDT |
| Read 1 Custom SeqB | GCC TGT CCG CGG AAG CAG TGG TAT CAA CGC AGA GTA C | 50 µM | Sigma Aldrich |
| Nextera_N701 | CAA GCA GAA GAC GGC ATA CGA GAT **TCG CCT TA**G TCT CGT GGG CTC GG | 10 µM | Sigma Aldrich |
| Nextera_N702 | CAA GCA GAA GAC GGC ATA CGA GAT **CTA GTA CG**G TCT CGT GGG CTC GG | 10 µM | Sigma Aldrich |
| Nextera_N703 | CAA GCA GAA GAC GGC ATA CGA GAT **TTC TGC CT**G TCT CGT GGG CTC GG | 10 µM | Sigma Aldrich |
